# Plant miRNA regulation is environmentally and developmentally-sensitive

**DOI:** 10.1101/136762

**Authors:** Patrick von Born, Ignacio Rubio-Somoza

## Abstract

Development and fitness of any organism rely on properly controlled gene expression. This is especially true for plants, as their development is determined by both internal and external cues. MicroRNAs (miRNAs) are embedded in the genetic cascades that integrate and translate those cues into developmental programs. miRNAs negatively regulate their target genes mainly post-transcriptionally through two co-existing mechanisms; mRNA cleavage and translational inhibition. It is unclear whether the efficiency of miRNA-guided regulation is generally influenced by factors like ambient temperature or developmental stage. Here we show that plant miRNA accumulation, as well as miRNAs’ mode of action can be temperature- and development-sensitive. Higher temperatures tend to induce a more pronounced accumulation of mature miRNAs. Both parameters have also an impact on the expression patterns of the core players involved in miRNA performance. We show that efficiency of miRNA-mediated gene silencing declines with age during vegetative development in a temperature-dependent manner. Co-existence of cleavage and translational inhibition was also found to be dependent on temperature and developmental stage. Therefore, each miRNA family specifically regulates their respective targets, while temperature and growth influence the performance of miRNA-dependent regulation in a more general way.

## Introduction

Control of gene expression is paramount for any organism in order to exist and transit through different developmental stages as well as interrelate with their surroundings during their life cycle. All layers of control of gene expression are tightly regulated, from chromatin state to protein post-translational modifications, including mRNA stability. Small RNAs (sRNAs) have emerged in the last decades as central elements embedded in those regulatory layers. sRNAs come in several flavors depending on the source of RNA used for their biogenesis (1). MicroRNAs (miRNAs) are a special class of sRNAs that mainly regulate the expression of their targets post-transcriptionally. miRNA-dependent regulation has evolved independently in at least six eukaryotic lineages, including land plants (2). Most of the current knowledge about plant miRNA biogenesis, action and function comes from studies in the model organism *Arabidopsis thaliana*. Primary miRNA transcripts (pri-miRNA) arise from the RNA polymerase II-dependent expression of independent transcriptional units. Their expression pattern is under the control of specific regulatory sequences as is the case for protein coding genes (3). Pri-miRNAs are processed by the microprocessor complex in mature miRNA duplexes ranging from 19 to 24 nt at the dicing bodies within the nuclei in a two-step enzymatic reaction (4). Proteins from the DICER family, mainly DICER-LIKE1 (DCL1;(5) are the core components of the microprocessor complex and are assisted by accessory proteins such as HYPONASTIC LEAVES1 (HYL1; (6) or DOUBLE-RNA BINDING PROTEIN 2 (DRB2;(7) SERRATE (SE; (8) and C- TERMINAL DOMAIN PHOSPHATASE-LIKE 1 (CPL1; (9). The resulting mature miRNA duplexes are subsequently protected from degradation through HUA-ENHANCER 1 (HEN1)-mediated methylation (5). Next, HASTY (HST; (10) participates in the transport of the stabilized miRNA duplexes to the cytoplasm where they are loaded into the RNA-Induced Silencing Complex (RISC). Proteins from the ARGONAUTE (AGO) family are the main executive components of the RISC complex. The Arabidopsis genome has 10 *AGO* genes of which AGO1 (11) and AGO10 are considered the main players in post-translational miRNA-mediated gene silencing (12). Once loaded into the RISC, one of the two duplex strands is degraded while the remaining one serves to scan the cytoplasm seeking for highly complementary mRNAs. miRNAs control the expression of their targets both by mRNA-target cleavage and translational inhibition (12). Beyond their existence, knowledge about the coexistence of both mechanisms in plants is scarce and suggests that the degree of their regulatory contribution might be cell-specific (13).

Plant miRNAs are involved in the regulation of a series of developmental and stress-related genetic programs (14, 15). Nevertheless, little is known about whether general miRNA biogenesis and action, or the efficiency of their regulation change during the course of development and/or as consequence of environmental changes. Initial attempts of dealing with such a gap relied on assaying changes of endogenous miRNAs (16, 17). A major drawback from those studies is that mature miRNAs are usually produced from polygenic families and their accumulation is driven by distinct chromatin modifications, promoter activity and pri-miR structure (18-20).

In order to circumvent such limitations and clearly discern how those parameters might influence miRNA performance, we used an artificial and sensitive miRNA reporter (9). Our results show that accumulation of mature plant miRNAs and the resulting regulation (mechanism and efficiency) of their targets are dependent on growth temperature and developmental stage. We also show that both factors affect the expression of several key players involved both in miRNA biogenesis and action.

The mechanisms of miRNA-mediated attenuation of protein expression have been harnessed to silence specific genes with artificial miRNAs (amiRs; (21, 22). Therefore, our findings are not just of relevance to understand miRNA regulation, but also instructive for the use of amiR-based gene silencing technology.

## Material and Methods

### Plant Material

Plants were grown on soil in long days (16 h light/8 h dark) under a mixture of cool and warm white fluorescent light at 16°C and 23°C and 65% humidity. *LUC* miRNA-activity reporter (9) and *rLUC* control in which synonymous point mutations were introduced to render the firefly luciferase miRNA-insensitive (23) have been previously described.

### RNA analyses

Total RNA was isolated as described in (24) using tissue pooled from 15 randomized individuals per sample and biological replicate.

Reverse transcription was performed with the RevertAid First Strand cDNA Synthesis Kit (Thermo Scientific) using 200 ng of total RNA previously treated with DNase I (Thermo Scientific) following the protocol described in (25).

PCRs were carried out in presence of SYBR Green (Invitrogen) and monitored with the CFX384 Real-Time PCR Detection-System (Bio-Rad) in two technical and two biological replicates. Biological replicates were treated as independent samples. Relative expression changes were calculated using 2^-ΔCt^ in all assays except in Fig. 3A where the 2^-ΔΔCt^ method was applied to normalize *LUC* mRNA levels to the ones of *rLUC*. Expression levels were normalized to *β-TUBULIN2* (At5g62690). Mature miRNA quantifications were performed by stem-loop RT-PCR as described (25).

For small RNA blots, 3 µg of total RNA were used and two biological replicates performed. All primers used are listed in Table S1.

### Protein assays

Proteins were isolated from the corresponding tissues from 15 randomized individuals per sample and biological replicate. After tissue homogenization, the resulting powder was resuspended in protein extraction buffer (PBS, Triton X-100 0.1%, Complete EDTA-free (Roche)). After centrifugation, 50 µl of protein were mixed with the same volume of Beetle-Juice (PJK) Firefly substrate. Luciferase activity from two biological replicates was measured in technical triplicates on a Centro LB 960 (Berthold Technologies) device. Protein concentration of two biological replicates was assessed using the Bradford protein assay kit (BioRad) in technical triplicates. From this, Luciferase activity per µg of protein was calculated and the average of both biological replicates was used for further analysis. Values were normalized to the ones from rLUC.

## Results

### Addressing developmental and environmental influence on miRNA-mediated regulation

RNA silencing has been described as an antiviral defense mechanism in both plants and invertebrates (26). Such defense mechanism is temperature sensitive with higher temperatures leading to increased production of virus-derived sRNAs (27). In order to study whether miRNA-mediated silencing is also under the influence of environment and/or development, we used an artificial miRNA reporter system that proved to be highly sensitive to perturbations in miRNA biogenesis and action (9). This reporter system relies on the expression of the Firefly luciferase gene (*LUC*) under the constitutive Cauliflower Mosaic Virus 35S promoter. Simultaneously, the expression of an artificial miRNA (*amiR- LUC*) driven by the very same promoter, specifically silences *LUC* expression. As control, we used a similar reporter system in which synonymous point mutations were introduced within the miRNA-complementary sequence in the *LUC* gene. Those point mutations rendered the LUC mRNA resistant (*rLUC*) to amiR-LUC regulation (23). Using that artificial approach has clear advantages compared to rely on endogenous miRNAs. Among those advantages, the production of both miRNA and target are controlled by the same promoter and can be related at all growth conditions and developmental stages to the proper control allowing a fine dissection of all steps of the regulation.

Arabidopsis plants carrying either the *LUC* or *rLUC* reporter systems were grown along at 16°C and 23°C. 16°C is closer to the temperatures Arabidopsis typically experiences in its normal habitats, while 23°C, despite being commonly used for Arabidopsis growth in controlled chambers, can be considered a stress temperature. Since the speed of Arabidopsis growth is temperature-dependent (28), we established discrete and equivalent time points to collect representative samples spanning the main developmental stages at both temperatures (Fig. 1A). Seedlings with the two first true leaves and leaves number 4 and 7 are representative of the transitions from juvenile to adult stages during vegetative development (Fig. 1B, (29). We also assessed inflorescences containing all closed buds (stages 1 to 12 (30) and pools of the three uppermost siliques after abscission of the senescent floral organs. Levels of the developmental timer miR156 were used to validate the equivalence of the samples collected at the two different growth conditions (31-33). As expected, miR156 accumulation declined as development progressed confirming that both sets of samples were developmentally equivalent (Fig. 1C). Slightly higher levels of miR156 observed in plants grown at lower temperatures are in line with former studies (16, 18).

**Fig. 1.**
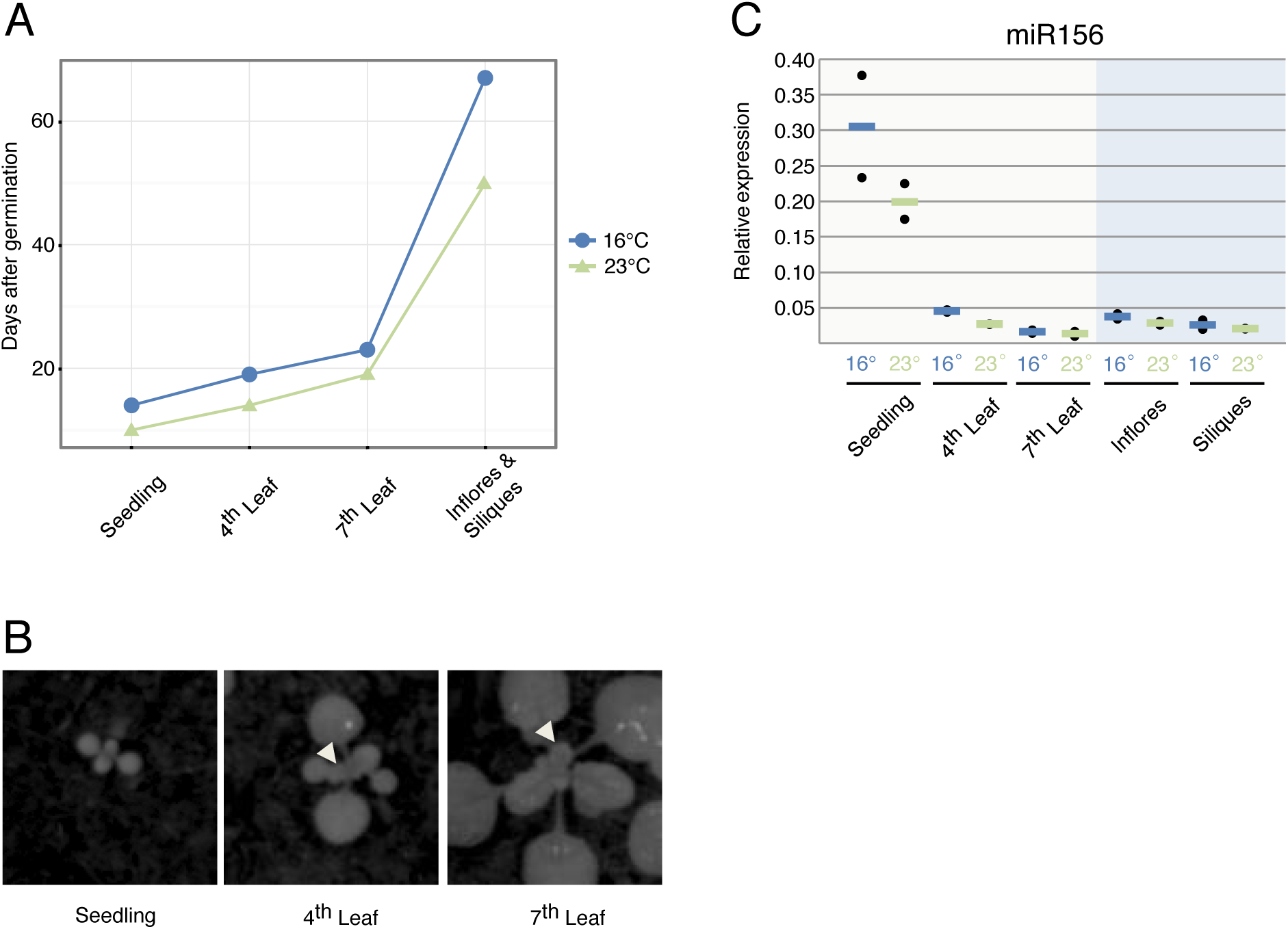
Addressing developmental and environmental influence on miRNA-mediated regulation via a Luciferase reporter system. (A) Discrete time points for tissue collection over Arabidopsis life cycle at 16°C (blue) and 23°C (green). (B) Representative pictures of the different leaf stages collected spanning vegetative development. Arrows point to the collected leaves. (C) Mature miR156 qRT-PCR to ensure that samples from both datasets (16°C and 23°C) were at equivalent developmental points. Black dots represent one biological replicate each, calculated from two technical replicates. Lines, (blue = 16°C, green = 23°C) represent the average between two biological replicates. “Inflores” stands for inflorescences.

### Mature miRNA accumulation has developmental and temperature-dependent components

To study accumulation of mature amiR-LUC, we assayed amiR levels by stem-loop qRT-PCR (Fig. 2A) and small RNA blots (Fig. 2B).

**Fig. 2.**
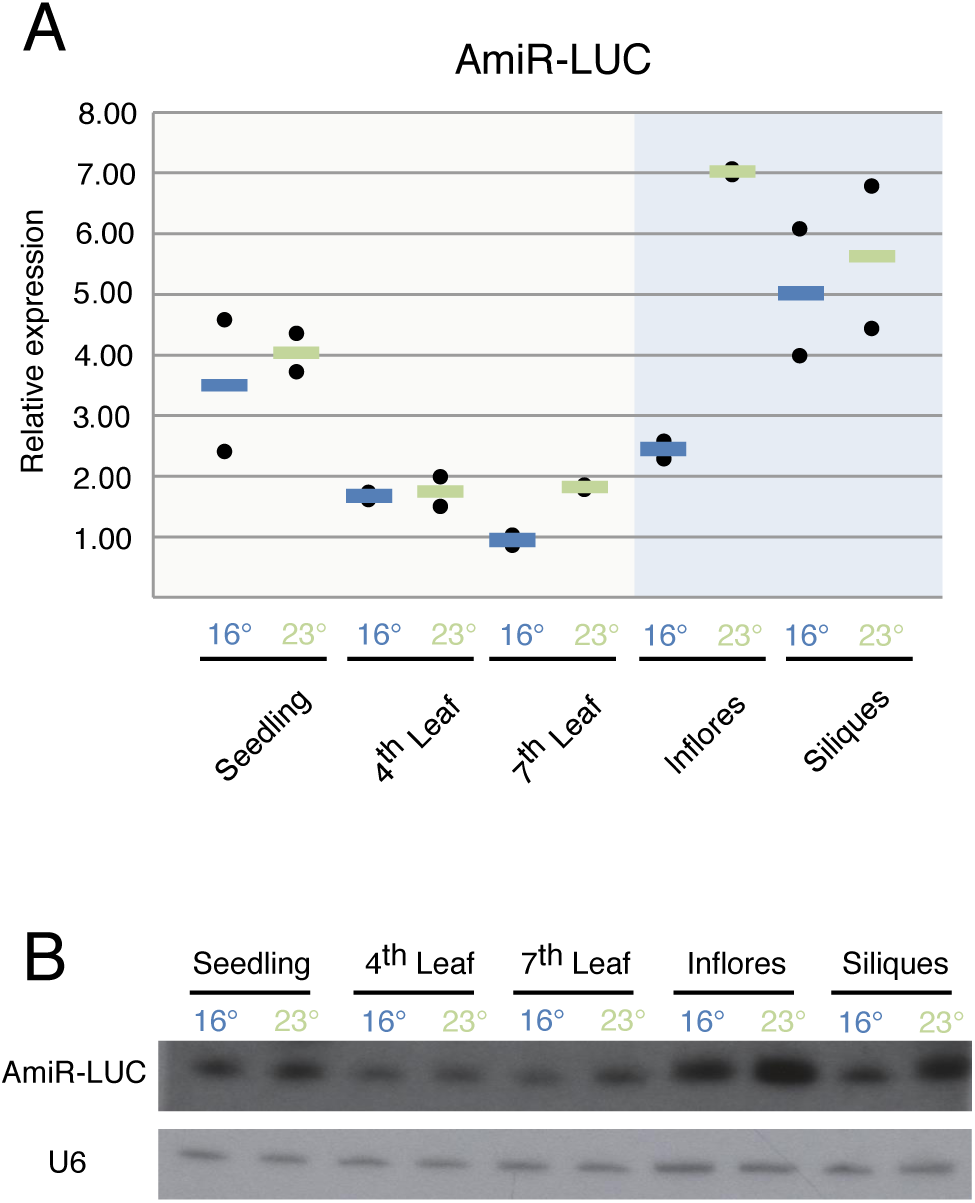
AmiR-LUC accumulation is developmentally and temperature-dependent. (A) Mature amiR-LUC accumulation assayed by qRT-PCR. Black dots represent one biological replicate each calculated from two technical replicates. Lines, (blue = 16°C, green = 23°C) represent the average between two biological replicates. “Inflores” stands for inflorescences. (B) Representative sRNA blot for amiR-LUC accumulation.

Independent of growth temperature, amiR-LUC accumulated to higher levels in seedlings than at later stages during vegetative development, *i.e.* leaves 4 and 7 (Fig. 2A, Fig. 2B). Moreover, amiR-LUC levels were higher in siliques and inflorescences at 23°C when compared to vegetative organs (Fig. 2A, Fig. 2B).

Higher temperature was found to increase amiR-LUC levels in late vegetative development (leaf 7) and especially in inflorescence (Fig. 2A, Fig. 2B).

Discrepancies found between the amiR-LUC levels determined either by qRT-PCR or small RNA blot in inflorescences and siliques might be explained by the intrinsic properties of both techniques (Fig. 2A, Fig. 2B). While stem-loop qRT-PCR monitors only the 21nt long species matching the designed amiR-LUC, small RNA blots can detect isoforms of different length and/or isoforms shifted by a few nucleotides (9).

A simple reason for miRNA accumulation being temperature and stage-dependent could be differential expression of factors involved in miRNA production. We therefore assayed whether the expression of core factors involved in miRNA biogenesis was regulated by development and/or growth temperature. We focused on the core executor DCL1 and in its assistants HYL1, DRB2, SERRATE and CPL1 (Fig. 3).

**Fig. 3.**
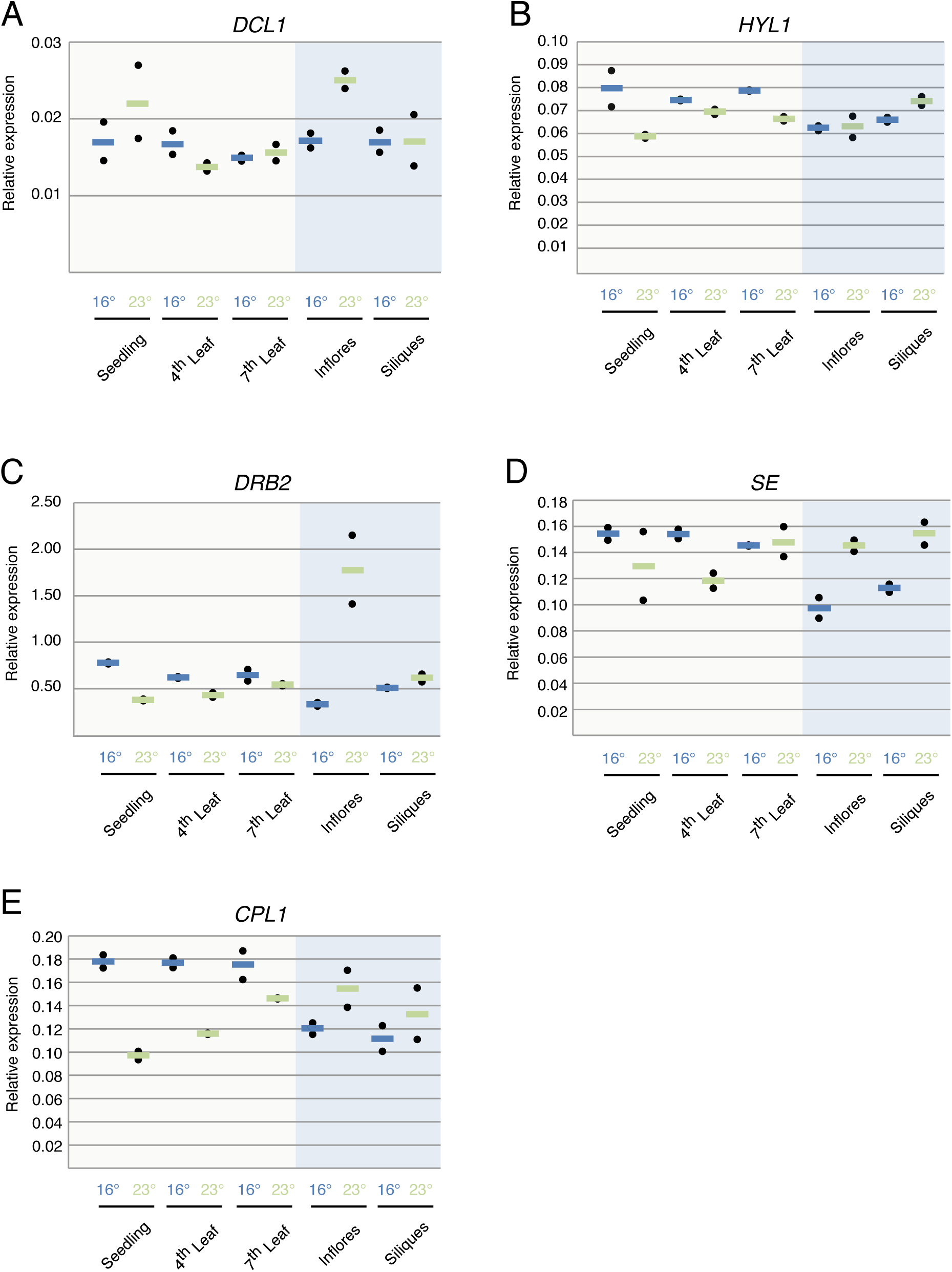
Effect of development and temperature on the expression of miRNA biogenesis factors. (A) DCL1. (B) HYL1. (C) DRB2. (D) SE. (E) CPL1. Black dots represent one biological replicate each calculated from two technical replicates. Lines, (blue = 16°C, green = 23°C) represent the average between two biological replicates. “Inflores” stands for inflorescences.

*DCL1* mRNA expression levels were similar across all samples, with the exception of a marked increase in inflorescences from plants grown at 23°C compared to their counterparts grown at lower temperature (Fig. 3A). *HYL1* and *DRB2* showed similar expression levels in all tested tissues with a common trend of lower expression in vegetative tissues at 23°C (Fig. 3B, Fig. 3C). Similar to DCL1, *DRB2* levels were increased in inflorescences from plants grown at 23°C (Fig. 3C).

*SE* was more highly expressed in vegetative than in reproductive organs in plants grown at 16°C (Fig. 3D). As for DCL1 and DRB2, SE expression levels in reproductive tissues were higher at 23°C.

Plants grown at 23°C presented the same trend of lower levels of *CPL1* expression in vegetative tissues that was also found for *HYL1* and *DRB2* (Fig. 3E). While its expression at 16°C did not change during vegetative development, it gradually increased in plants grown at 23°C reaching highest values in leaf 7 (Fig. 3E).

Collectively, our results show dynamic and heterogeneous expression profiles of different members of the core miRNA biogenesis machinery. We observed little correlation between these patterns and the accumulation of mature amiR-LUC across the different samples with the only exception of inflorescences from plants grown at 23°C. When compared to plants grown at 16°C, higher levels of amiR-LUC were paralleled by higher levels of *DCL1, DRB2, SE* and *CPL1*. Further, the slightly lower levels of expression for *HYL1* coincide with *DRB2* expression levels during vegetative development at 23°C.

It is noteworthy that for most of the miRNA biogenesis players, we observed a general tendency to higher expression levels in vegetative organs at 16°C than in plants grown at 23°C, while the opposite was true in reproductive organs.

### Efficiency and mode of action of miRNA-mediated regulation is temperature dependent

Once we had established that development and temperature affect the accumulation of mature miRNAs, we sought to explore whether miRNA-mediated gene silencing was also developmentally and environmentally regulated.

We firstly assayed the contribution of target cleavage regulation in response to different growth temperatures and across development. *LUC* mRNA levels were assayed in the same samples used for qRT-PCR and with primers flanking the miRNA-targeted sequence. *LUC* levels were reduced by 60 to 85% when compared to *rLUC* depending on tissue and growth conditions (Fig. 4A). We found that generally, higher levels of mature amiR-LUC (Fig. 2) lead to lower levels of *LUC* transcripts (Fig. 4A). Thus, seedlings and inflorescences from plants grown at 16°C presented higher levels of *LUC* transcripts than their counterparts grown at 23° (Fig. 4A). During vegetative development, we observed an increase in *LUC* mRNA levels between seedlings and leaf 4 both at 16°C and 23°C (Fig. 4A).

**Fig. 4.**
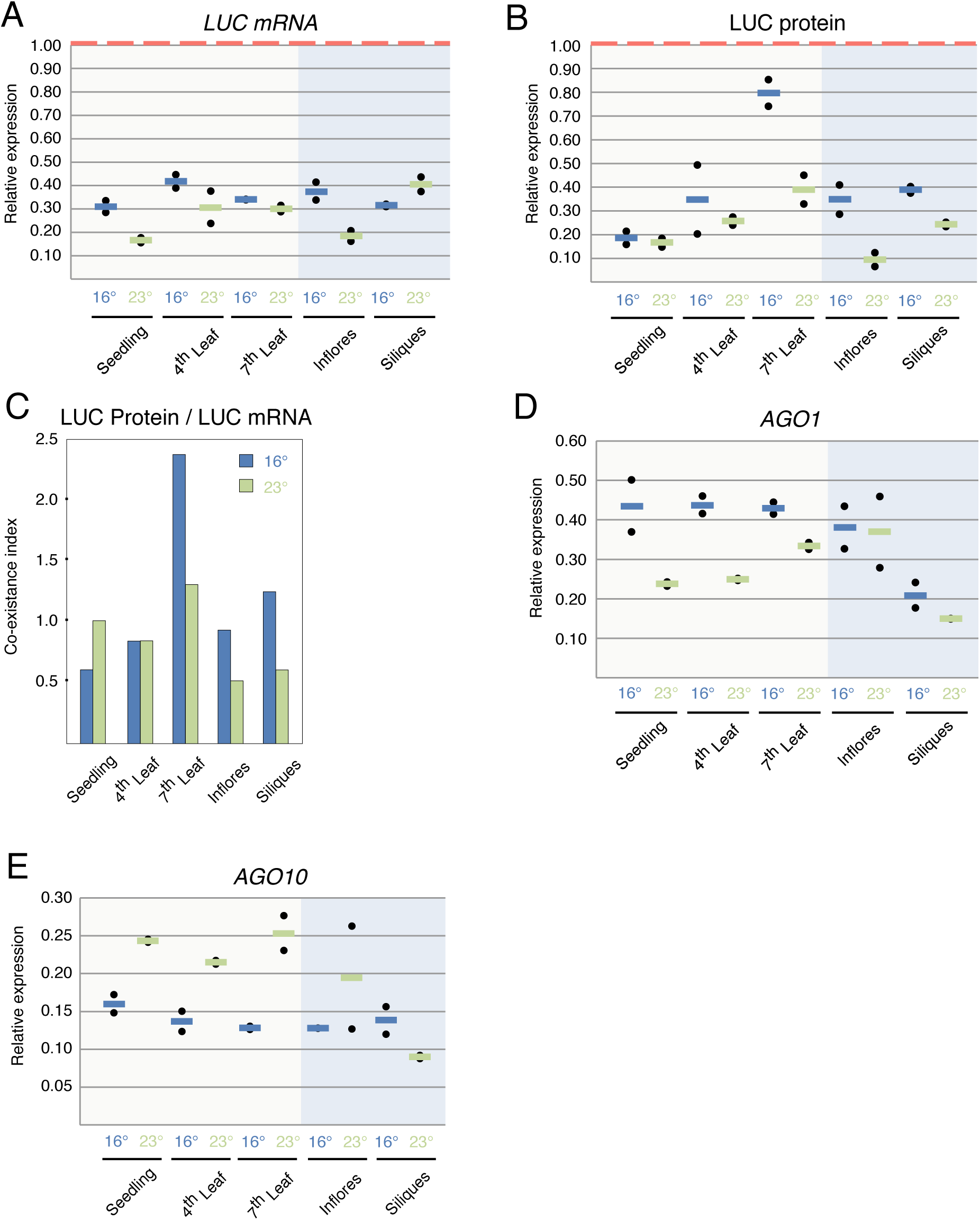
miRNA mode of action is developmentally and temperature-dependent. (A) LUC mRNA expression levels assayed by qRT-PCR normalized to LUC mRNA in rLUC control plants (red dotted line). Lines, (blue = 16°C, green = 23°C) represent the average between two biological replicates. (B) LUC protein levels. Black dots represent one biological replicate each calculated from two technical replicates. Lines, (blue = 16°C, green = 23°C) represent the average between two biological replicates normalized to LUC protein levels in rLUC control plants (red dotted line). (C) Coexistence index is the ratio of average protein levels by average mRNA levels from each sample and condition. (D) AGO1 expression levels assayed by qRT-PCR. Black dots represent one biological replicate each calculated from two technical replicates. Lines, (blue = 16°C, green = 23°C) represent the average between two biological replicates. (E) AGO10 expression levels assayed by qRT-PCR. Black dots represent one biological replicate each calculated from two technical replicates. Lines, (blue = 16°C, green = 23°C) represent the average between two biological replicates. (A-E) “Inflores” stands for inflorescences.

To assess the contribution of translational inhibition we inferred the levels of LUC protein by measuring LUC activity in protein extracts from samples collected at the same time as the ones used for expression assays (Fig. 4B). We observed that during vegetative development LUC levels gradually increased, although to a different extent, both at 16°C and 23°C (Fig. 4B). We also found that the ultimate effect of miRNA-dependent regulation over the production of functional targeted-protein was temperature dependent. LUC protein levels were clearly higher at 16°C than the ones found in samples from plants grown at 23°C in leaf 7, inflorescences and siliques (Fig. 4B).

Next, we studied whether the differential contribution of both regulatory mechanisms was developmentally and/or environmentally determined. We reasoned that translational inhibition mechanisms would lead to a further reduction of LUC protein levels when compared to mRNA levels. Therefore, we created a Coexistence index, representing the ratio between LUC protein levels and LUC mRNA levels. Values higher than 1 indicated a low contribution from translational inhibition to miRNA-dependent regulation, while the opposite was true for values smaller than 1. As seen in Fig. 4C, the translational inhibition mechanism was gradually less effective during vegetative development at 16°C. We observed the same tendency in leaf 7 from plants grown at 23°C when compared with earlier stages of development (seedlings and leaf 4). In inflorescences and siliques, translational inhibition was more potent at 23°C when compared to 16°C.

The two main effectors within miRNA-loaded RISC complexes are AGO1 and AGO10. Both proteins have redundant but also specific roles in miRNA-mediated gene silencing (34). Thus, it has been suggested that AGO10 has a more prominent role on translational inhibition (34) despite evidence that it is also able to cleave its mRNA targets (35). To ascertain whether developmental and environmentally-dependent changes on the coexistence index correlated with variations on their expression, we analyzed both *AGO1* and *AGO10* profiles by qRT-PCR (Fig. 4D; Fig. 4E). Interestingly, during vegetative development both *AGO1* and *AGO10* presented an opposite expression pattern. *AGO1* expression was consistently higher at 16°C while the opposite was true for *AGO10*. Nevertheless, we did not observe any correlation between *AGO1* and *AGO10* expression patterns and the differences in co-existence index.

Altogether, these results show that developmental as well as environmental components influence both miRNA regulation and the balance between cleavage and translational inhibition mechanisms of gene silencing.

## Discussion

Our findings show that plant miRNA performance (accumulation, efficiency and co-existence of target cleavage and translational inhibition) is influenced by both development and environment. Our results support that the expression of several central players in miRNA performance also depends on development and temperature in which plants are grown.

The view of the different pathways involved in sRNA production and action was initially rather simplistic and static (36). It was generally assumed that molecular players devoted to generate each type of sRNA were ubiquitously expressed and, therefore, the main layer of control on sRNA-mediated regulation was orchestrated by the expression patterns of the RNA from which they originated. We are currently starting to appreciate that this might be a more dynamic process (37). Our results support a more dynamic scenario in which the expression of molecular players and mechanisms involved in miRNA-mediated gene silencing are developmentally and environmentally-sensitive.

Although siRNA biogenesis in plants has been reported to be temperature sensitive, with siRNA levels correlating with growth temperature, mature miRNA accumulation has been thought to be largely temperature insensitive (27, 38, 39). In contrast to studies where whole plants were assayed, our study dissects the temperature effect using discrete samples that encompass the different developmental stages during vegetative and reproductive development. Our analysis shows that amiR-LUC accumulation is temperature-responsive in leaves produced at later stages (leaf 7) when compared to an early phase of vegetative development (leaf 4). That positive temperature effect on amiR- LUC levels is more dramatic in reproductive development with a greater accumulation in inflorescences grown at 23°C (Fig. 2A; Fig. 2B). Such increased accumulation is likely to be a consequence of the higher expression of the central miRNA biogenesis factor *DCL1* (Fig. 3A) and its assistants *DRB2* (Fig. 3C) and *SE* (Fig. 3D). Among the factors involved in miRNA biogenesis, temperature had a clear effect on the expression of *SE* and *CPL1* which are additionally involved in more general processes such as mRNA splicing, which seems to be also temperature dependent (40-43).

miRNA-mediated gene silencing relies on two mechanisms that are thought to coexist, target cleavage and translational inhibition (12). Nevertheless, beyond their existence little is known about their individual contribution to target gene silencing in plants. In mammals, miRNA-mediated regulation occurs mainly through target degradation regardless of cell type, growth conditions or translational state (44, 45). Initial work shows that in plants the contribution of both mechanisms is cell-specific in pollen (13). It is also unknown whether environmental conditions can influence plant miRNA efficiency and their mode of action.

Our results reveal that the efficiency of miRNA regulation decays with age in Arabidopsis (Fig. 4C; leaf 4 versus leaf 7) with this decrease being more evident at low temperatures. Likewise, the contribution of translational inhibition also decreases with age showing a similar discrepancy between mRNA and protein levels to the one found in mutant plants impaired in that mechanism of gene silencing (12). In contrast, the contribution from this mechanism increases with temperature during reproductive development (Fig. 4C). Although we found an opposite effect of temperature on the expression of the two main silencing effectors, *AGO1* and *AGO10*, we did not find any correlation between their levels and the difference in miRNA efficiency or mode of action. It has been suggested that miR168-loaded AGO10 negatively regulates AGO1 protein levels without affecting its mRNA levels (34). Our results suggest that AGO10 might also affect *AGO1* mRNA levels.

According to our results, miRNA regulation is more efficient at higher temperatures. Therefore, it is tempting to hypothesize miR168 AGO10-loaded contributing to *AGO1* mRNA degradation at 23°C. In support of such hypothesis, it has been described that AGO10 has slicing activity (35). This point deserves further investigation.

It has been recently suggested that the DCL1 partner proteins HYL1 and DRB2 determine whether a miRNA triggers cleavage or translational repression of its targeted mRNAs (46). Thus, while HYL1-mediated miRNA production contributes to degradation of the targeted mRNA, DRB2-dependent miRNA biogenesis triggers translational inhibition. Despite the observed changes in the coexistence between both regulatory mechanisms over development, we could only correlate higher levels of *DRB2* expression to a more pronounced contribution through translational inhibition in inflorescences grown at 23°C when compared to lower temperature (Fig. 2C, Fig. 4C). Absence of correlation suggests that additional players and/or posttranslational modifications of the already known ones might determine the mechanism through which miRNAs regulate the expression of their targets (9, 47). Therefore, in contrast to animals, plant miRNA performance depends on the cell-type, developmental stage and growth conditions.

Plants compromised in essential components of the miRNA machinery, such as DCL1 and AGO1 (48, 49), are usually sterile when grown at 23°C. Nevertheless, a partial restoration of fertility is found when those plants are grown at lower temperatures. According to our results, miRNA regulation efficiency in inflorescences is lower at 16°C when compared to plants grown at higher temperatures. Consequently, miRNA gene silencing might play a minor role in the general regulation of gene expression at low temperatures in inflorescence thereby explaining fertility restoration in these growth conditions.

Finally, our results are informative for the use of artificial miRNAs to downregulate endogenous genes at late stages of development or as part of crop protection strategies.

## Acknowledgments

We greatly acknowledge the support from Detlef Weigel since majority substantial part of this work was conducted at the MPI for Developmental Biology. We thank Silvio Collani, Catia Igreja and Elisa Izaurralde for support with establishing the luciferase quantification assay, and Patricia Lang, Sascha Laubinger, Peter Brodersen, Pablo Manavella and members of the MoRE Lab for critical reading of the manuscript.

## Contributions

I.R-S. designed the research, performed experiments, analyzed the data, and wrote the manuscript. P.v.B. performed experiments, analyzed data and helped in manuscript drafting.

